# A robust phylogenetic framework for ophiostomatoid fungi: orders Ophiostomatales and Microascales (Sordariomycetes, Ascomycota)

**DOI:** 10.64898/2026.01.27.701938

**Authors:** Theo Llewellyn, Alfried P. Vogler

## Abstract

Ophiostomatoids are an ecological group of microfungi that commonly associate with bark and ambrosia beetles. As well as being insect symbionts, they play significant ecological roles as plant pathogens, and include species responsible for major forest tree diseases. Despite their ecological similarities, ophiostomatoids are distributed across two quite distantly related orders, the Microascales and Ophiostomatales. Historically, these fungi were considered a single natural group; however, molecular studies have revealed their independent origins and convergent ecological strategies. Previous phylogenetic studies of these fungi have typically focused on resolving taxonomic issues or understanding individual lifestyles, such as beetle-cultivated ambrosia lineages or vascular wilt pathogens. As a result, we lack a comprehensive phylogenetic framework that integrates dense species-level sampling with ecological data across both orders. Such frameworks are essential for understanding the broader phylogenetic and ecological context in which key fungal lifestyles have evolved. Here, we assembled and analysed all available sequence data for the Microascales and Ophiostomatales from seven widely used fungal marker loci to reconstruct a densely sampled phylogeny for each order. We evaluated locus performance and showed that whilst individual loci fail to resolve many taxa, concatenated datasets produce robust, well-supported topologies consistent with published genomic studies. By mapping ecological traits onto these trees, we show that lifestyle diversity and beetle associations are much more variable in the Microascales than in the Ophiostomatales, despite comparable species richness. Presenting both orders together provides a unique comparative perspective on the ecology and evolution of ophiostomatoids. As metabarcoding datasets of ophiostomatoids become increasingly common, this integrative framework can offer a valuable resource for environmental sequence identification and investigating fungal lifestyle switches, which in turn can support future biodiversity and ecology studies.

## Introduction

Ophiostomatoids are a polyphyletic group of fungi from the Ophiostomatales and Microascales orders (Sordariomycetes, Ascomycota). Despite being quite distantly related within the Sordariomycetes (Georg Hausner, Reid and Klassen, 1993; G. Hausner, Reid and Klassen, 1993a, 1993a; Spatafora and Blackwell, 1994; Li *et al*., 2021), these fungi share key morphological traits related to insect dispersal, such as flask-shaped ascomata and sticky spore droplets (Luttrell, 1951; Benny, Benny and Kimbrough, 1980; von Arx and van der Walt, 1988). They are commonly associated with bark and ambrosia beetles (Scolytinae and Platypodinae, Curculionidae), forming a range of symbiotic interactions from facultative and commensal dispersal to obligate reciprocal mutualism (Hulcr and Stelinski, 2017; Biedermann and Vega, 2020). The most famous of these cause major plant diseases like Dutch elm disease and laurel wilt (Brasier, 1991; Harrington, Fraedrich and Aghayeva, 2008). Some associations also represent exceedingly rare cases of beetle-mediated “fungal agriculture” (Farrell *et al*., 2001).

Although ophiostomatoids are distributed across two distinct orders, their shared morphology led taxonomists to consider them a single natural group for many years (Nannfeldt, 1932; Luttrell, 1951). Only as late as 1980, Benny & Kimbrough (1980) recognised the Microascaceae and Ophiostomataceae as members of different orders based on conidia and ascospore structure: describing the new order Ophiostomatales, and redefining the Microascales to contain the Microascaceae, Chadefaudilleaceae, and Pithoascaceae. At that time, the Ophiostomataceae contained four genera: *Ophiostoma, Ceratocystiopsis*, *Sphaeronaemella*, and *Ceratocystis*. Shortly after, Upadhyay (1981) synonymised *Ophiostoma*, *Sphaeronaemella*, *Grosmannia* and *Europhium* with *Ceratocystis* based on similar long-necked ascocarp producing sticky, sheathed spores, making *Ceratocystis* the type genus of Ophiostomataceae.

Subsequent molecular studies revealed that the Ophiostomatales genera *Ophiostoma* and *Ceratocystis* were in fact quite distantly related, and *Ceratocystis* was moved out of the Ophiostomatales and back into the Microascales (Georg Hausner, Reid and Klassen, 1993; G. Hausner, Reid and Klassen, 1993a, 1993b; Spatafora and Blackwell, 1994). Since then, both orders have undergone considerable taxonomic revision. Most Ophiostomatales changes have focused on genus description and delimitations (Zipfel *et al*., 2006; De Beer, Seifert and Wingfield, 2013; de Beer, Duong and Wingfield, 2016; de Beer *et al*., 2016; van der Linde *et al*., 2016; Bateman *et al*., 2017; Poinar and Vega, 2018; Nel *et al*., 2021). Recently, de Beer *et al*. (2022) produced a full genus-level revision for the Ophiostomatales based on sequences from four loci and over 200 species, supported by a genome-level phylogenetic backbone.

Similarly, the Microascales have seen many genus-level revisions (De Beer, Seifert and Wingfield, 2013; de Beer *et al*., 2014; Mayers *et al*., 2015; Nel *et al*., 2018; Mayers, Harrington, Masuya, *et al*., 2020) but also significant changes at the family level, expanding from four families to eight in the past two decades (Cannon and Kirk, 2007; Kirk *et al*., 2008; Réblová, Gams and Seifert, 2011; Wijayawardene *et al*., 2022; Bao *et al*., 2023).

Both orders have unresolved phylogenetic issues, for example, whether to split the single Ophiostomatales family Ophiostomataceae into smaller, more morphologically consistent families (de Beer *et al*., 2022), and whether the Microascales genera *Tubakiella* and *Nautosphaeria* could represent a new family for the order (Sakayaroj, Pang and Jones, 2011; Hyde *et al*., 2020; Bao *et al*., 2023). Additionally, the placement of many genera remains uncertain in both the Microascales (De Beer, Seifert and Wingfield, 2013; Bao *et al*., 2023) and Ophiostomatales despite the use of genomic data (Vanderpool, Bracewell and McCutcheon, 2018; Nel *et al*., 2021).

To resolve taxonomic issues, we need well-sampled, robust phylogenetic trees. However, habitat or lifestyle-specific sampling has resulted in uneven taxon coverage in many phylogenetic datasets. Although Microascales and Ophiostomatales are best known for their beetle associations, both orders contain a huge diversity of other ecological lifestyles, from wood-decaying saprotrophs specialised on marine driftwood (Halosphaeriaceae, Microascales) to globally distributed human pathogens like *Sporothrix schenckii* (Ophiostomatales) (López-Romero *et al*., 2011; Sakayaroj, Pang and Jones, 2011). This suggests that beetle association is highly homoplasic across ophiostomatoids, highlighting them as an intriguing case study of convergent lifestyle evolution that robust phylogenies could help us explore in greater detail.

To provide a more comprehensive basis for resolving phylogenetic issues and testing such evolutionary hypotheses, we compiled all publicly available sequence data for the Microascales and Ophiostomatales across seven widely used fungal nuclear markers to reconstruct densely sampled phylogenies for each order. We coupled these trees with ecological lifestyle and beetle association data to explore patterns of fungal lifestyle diversity both within and between orders. By presenting the two phylogenies together, we provide the research community with a consolidated, accessible framework that can support future work on ophiostomatoid diversification, evolutionary ecology, and symbiotic interactions.

## Methods

### Sequence Data Curation and Taxon Sampling

We compiled sequence data for representatives of the Microascales and Ophiostomatales, along with their closest related outgroups, based on the genome tree of the kingdom Fungi (Li *et al*., 2021). For Ophiostomatales, Magnaporthales was the outgroup (genera *Pseudopyricularia*, *Proxipyriculria*, *Bambusicularia*, *Magnaporthe*, *Ophioceras*, *Gaeumannomycella*) and for Microascales, we used Glomerellales (*Colletotrichum* and *Verticillium*). Seven gene regions were targeted: three from the nuclear rRNA operon gene cluster, namely the Internal Transcribed Spacer (ITS), the Small Subunit (SSU/16S) and the Large Subunit (LSU/28S) rDNA, as well as protein-coding loci β-tubulin, Elongation Factor-1α (EF1a), DNA-directed RNA polymerase II largest subunit gene (RPB1), and DNA-directed RNA polymerase II second largest subunit gene (RPB2).

The following steps were performed for Microascales and Ophiostomatales separately. For in-group sampling, we downloaded all sequences on the NCBI nucleotide database under the taxon IDs for Microascales (taxid:5592) and Ophiostomatales (taxid:5151). We downloaded representative outgroup sequences from the 6-gene Fungi reference set v3 in the Tree-Based Alignment Selector (T-BAS) toolkit (Carbone *et al*., 2019). We checked for and removed duplicate sequence headers and only kept sequences between 200 and 5000 base pairs. We then used two custom R scripts from the phylogenie package to separate the downloaded sequences into the seven loci (2_Bind_and_clean_seqs.R and 3_Bait_and_pattern_sort.R). These scripts aligned sequences to reference sequences for each of the seven loci and searched for all potential locus names in the fasta headers as an additional source of information for sequence sorting.

For each of the seven loci, we sorted sequences by name and retained the longest or most complete sequence when multiple sequences were available for a single strain. Given our aim of producing species-level reference phylogenies for molecular systematics and phylogenetic placement, we removed any unnamed or environmental sequences. Identical sequences were only retained if the FASTA header clearly indicated they were from different, well-delimited taxa according to the literature.

### Sequence Processing and Alignment

Raw FASTA headers were standardised to TAXON_NAME_STRAIN using command-line sed functions to ensure consistency across loci. Sequences for each gene were aligned using MAFFT v7 with the ‘—auto’ strategy (Katoh, 2002). For the Microascales β-tubulin alignment, exon-only sequences required special handling, as standard MAFFT failed to correctly align the different subunits; instead, these shorter fragments were aligned against a complete *Colletotrichum* CDS reference (accession: PQ530035.1_cds_XKC23079.1_1) using MAFFT’s --add function.

Alignments were manually inspected and curated in AliView and duplicate or mislabelled sequences were removed. We also standardised strain IDs across loci. As ITS, SSU, and LSU regions are often sequenced and uploaded as a single DNA fragment, we used ITSx v1.1.3 (Bengtsson-Palme *et al*., 2013) to predict the start and end positions of each gene and trim any flanking regions. We identified poorly aligned sequences in each locus alignment and used Basic Local Alignment Search Tool (BLAST) (Camacho *et al*., 2009) searches to find and remove any misannotated or contaminant sequences. After cleaning the alignments, gaps were removed, and the sequences were realigned.

For each locus, we used MEGA11 (Stecher, Tamura and Kumar, 2020; Tamura, Stecher and Kumar, 2021) to calculate the GC%, AT skew, and the average pairwise distance using the Kimura 2-parameter (K2P) model and 100 bootstrap replicates (Kimura, 1980). For K2P distances, all ambiguous positions were removed for each sequence pair (pairwise deletion option). Additionally, we performed Chi-square tests of homogeneity of base frequencies in the command line implementation of PAUP v4.0a to see whether base frequency differed across taxa in each alignment (Swofford, 2003).

### Phylogenetic Tree Inference

Individual locus alignments were concatenated using AMAS (Borowiec, 2016). Different loci from the same strain were determined by strain IDs or known fungal culture collection accession codes, such as CBS accessions in the FASTA headers. For ambiguous cases, we checked the original publications to link sequence IDs to strain IDs. Using both strain IDs and culture codes allowed us to connect sequences from different studies that used the same original sample material. Partitioning schemes were generated in AMAS. As we were aiming for species-level resolution in our trees, we generally removed duplicates of the same latin binomial and retained only the concatenated sequence covering the highest number of loci. If two concatenated sequences covered equal numbers but different loci, both sequences were retained. Where possible, we also tried to retain concatenated sequences containing the universal fungal barcode region ITS, to make our trees and alignments more usable for future studies using phylogenetic placement.

Maximum likelihood phylogenetic trees searches were performed with IQ-TREE v2 (Minh *et al*., 2020). For species tree analyses, we employed ModelFinder with each locus in a separate partition but allowing for partition merging (-m MFP+MERGE) (Kalyaanamoorthy *et al*., 2017). We evaluated nodal support using SH-aLRT (1,000 replicates), ultrafast bootstrap (1,000 replicates), aBayes, and local bootstrap analyses. Individual gene trees were also estimated with IQ-TREE using the partitioned dataset (Chernomor, Von Haeseler and Minh, 2016; Hoang *et al*., 2018). Using IQ-TREE, we also calculated gene concordance factors (gCF), which measure the proportion of gene trees supporting each branch in the concatenated species tree, and site concordance factors (sCF), which measure the proportion of sites in the concatenated alignment that support each species branch (Minh, Hahn and Lanfear, 2020). For each gene, we also calculated the normalised quartet score using ASTRAL-III (Zhang *et al*., 2018). This represents the proportion of quartets in each gene tree that are observed in the species tree.

We identified long branches or anomalous placements suggestive of misidentification, misalignment, or sequencing artefacts. Sequences flagged as problematic were realigned where possible or removed from the dataset. Alignments and trees were reanalysed iteratively until no major anomalies remained. Trees were rooted using pxrr from the phyx package (Brown, Walker and Smith, 2017). For the final trees, we calculated the taxonomic retention index (tRI) to determine the level of taxonomic monophyly using the phylofuncs.R script from the phylostuff repository (https://github.com/tjcreedy/phylostuff).

### Fungal lifestyle data curation

Given the diversity of fungal lifestyles in the Ophiostomatales and Microascales, we mapped fungal lifestyle data from the FungalTraits database (Põlme *et al*., 2020) onto the concatenated maximum likelihood phylogenies to observe how lifestyles are distributed across the trees. Lifestyles were assigned to clades at the genus level, given the lack of species-level resolution in FungalTraits. We also performed a literature search to identify which genera or species complexes are associated with bark and ambrosia beetles.

## Results

### Microascales

We downloaded 33,616 Microascales sequences, 16,961 of which could be confidently sorted into one of the seven loci. After standardising names, removing identical sequences from the same taxon and identifying mislabelled or misidentified sequences, 14,536 remained. For each locus, we selected the longest sequence for each taxon and only kept sequences identified to species level, resulting in 1725 total sequences. After initial tree building, 316 sequences were further identified as misidentified, mislabelled or contaminants, leaving 1409 sequences across all seven genes. Concatenating loci resulted in a final Microascales alignment with 597 concatenated sequences and 12814 sites, of which 4180 were informative. The final alignment covered all Microascales families except two small families, Cornuvesicaceae and Triadelphiaceae. Supplementary Table 1 shows the taxa in the concatenated alignment and the NCBI accession codes for each sequence.

To evaluate the contribution of individual loci to phylogenetic inference, we compared locus alignment statistics and gene-wise clade support against the concatenated species tree. The number of available sequences varied widely among loci, from 414 for ITS to only 14 for RPB2 (Table 1). The number of informative sites was highest for β-tubulin (1085/3851) and lowest for SSU (135/1748). AT skew varied considerably between loci (Figure 1a), whilst GC content remained relatively stable (Figure 1b). Mean K2P distance was highest in RPB1 and RPB2 and particularly low in SSU. Nucleotide base frequencies differed significantly between taxa for ITS, LSU, EF1a and RPB2 (Chi-2 homogeneity of variance *p* < 0.05) (Figure 4f).

**Figure 1:**
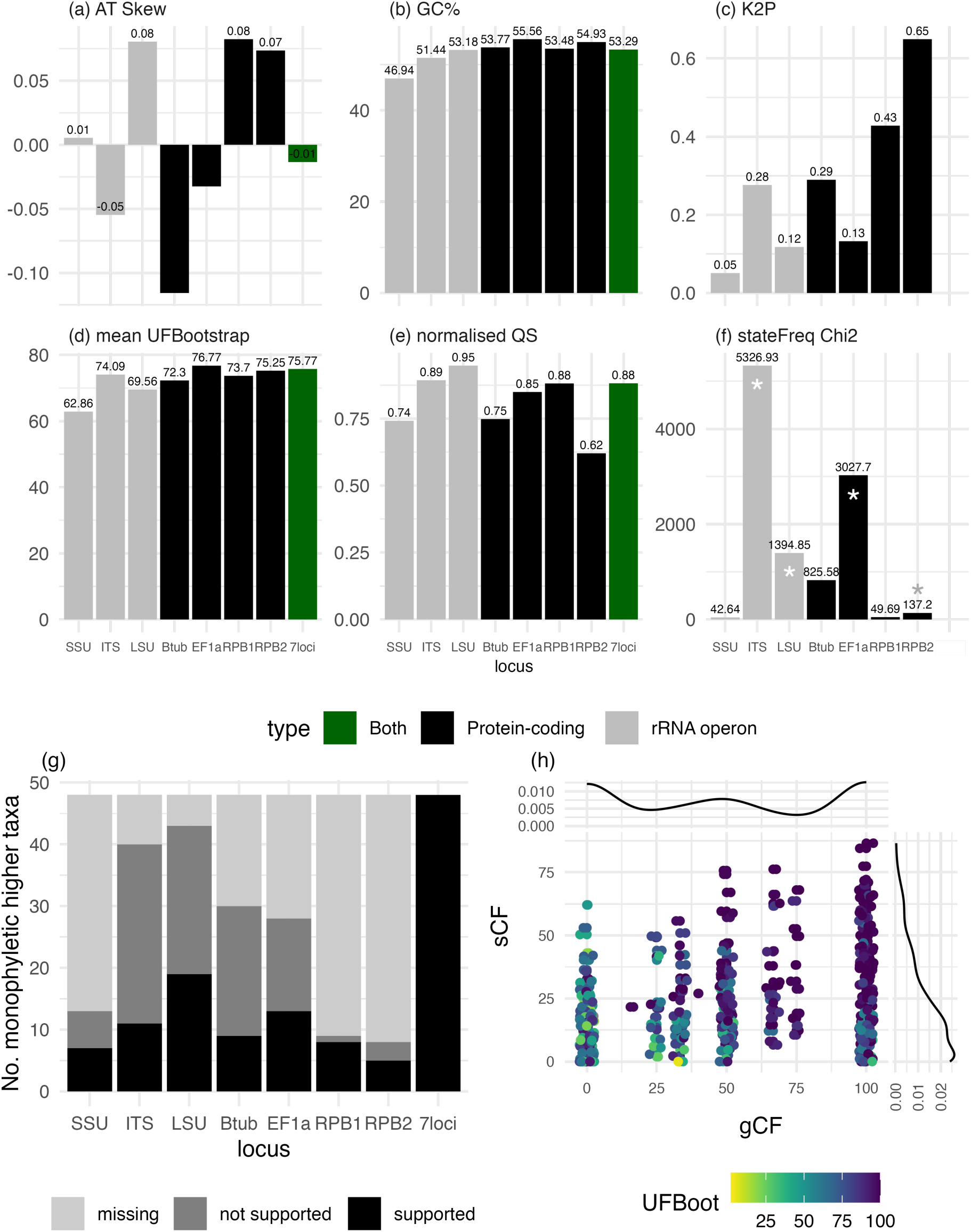
Plots comparing summary statistics of sequeunce alignments and phylogenetic trees for each locus and the concatenated 7-locus dataset. (a) Alignment AT Skew, (b) alignment GC%, (c) Kimura 2-parameter (K2P) pairwise distance, (d) mean UltraFast Bootstrap support, (e) normalised Quartet Score (QS), (f) nucleotide state frequency Chi-square with asterisk showing significance (*p* < 0.05), (g) number of higher taxa recovered as monophyletic (black), not monophyletic (dark grey), and missing (light grey), (h) scatter plot showing the relationship between IQTree gene Concordance Factor (gCF), site Concordance Factor (sCF), and UFBootstrap. Each point represents one node in the concatenated species tree. Histograms show the distribution of concordance factors.

**Table 1:** Alignment summary statistics for the Microascales 7-locus concatenated alignment and individual locus alignments.

| Locus | Sequences | Sites | Unique | Informative | Invariant |
| --- | --- | --- | --- | --- | --- |
| Concatenated | 596 | 12814 | 7229 | 4180 | 6835 |
| B-tubulin | 302 | 3851 | 2008 | 1085 | 2276 |
| EF1a | 270 | 2152 | 1424 | 912 | 1025 |
| ITS | 414 | 1201 | 1016 | 703 | 312 |
| LSU | 347 | 1783 | 1068 | 471 | 956 |
| SSU | 38 | 1748 | 425 | 135 | 1471 |
| RPB1 | 23 | 820 | 545 | 418 | 335 |
| RPB2 | 14 | 1259 | 756 | 456 | 460 |

When comparing locus trees, mean bootstrap support was noticeably lower in the SSU tree than in the other loci (Figure 4d). In general, locus trees agreed with the species tree (normalised Quartet Score = 0.88). However, individual locus scores varied when compared to the species tree, with LSU showing the highest QS of 0.95 and RPB2 the lowest at 0.62 (Figure 1e). We then investigated which described higher taxa seen in the species tree were also recovered in locus trees. Again, LSU supported the highest number of known higher taxa and RPB2 the lowest (Figure 1g). Supplementary Table 2 shows the full breakdown of which genes support which genera and families. For most nodes in the species tree, the site concordance factor (sCF) was very low (Figure 1h, Supplementary Figure 1). Gene concordance factors (gCF) showed a split distribution, with many receiving 100% gCF whilst a similar number received very low or even 0% gCF (Figure 1h).

In the maximum likelihood species tree based on the concatenated gene data, the order Microascales was strongly supported as monophyletic, subdivided into two clades: Ceratocystidaceae + Gondwanamycetaceae and Microascaceae + Graphiaceae + Halosphaeriaceae (Figure 2). Within the monophyletic Ceratocystidaceae, the genera *Ceratocystis*, *Davidsoniella*, *Endoconidiophora*, *Bretziella*, *Wolfgangiella*, *Toshionella*, *Ambrosiella*, *Huntiella*, *Berkeleyomyces*, and *Thielaviopsis* were supported monophyletic groups. *Chalaropsis* was not monophyletic, with the ovoidea group falling in a separate clade. Gondwanamycetaceae and both its genera, *Knoxdaviesia* and *Custingophora*, were strongly supported. The Graphiaceae family was also well-supported. The average taxonomic retention index (tRI) for the Microascales species tree was 0.6137.

**Figure 2:**
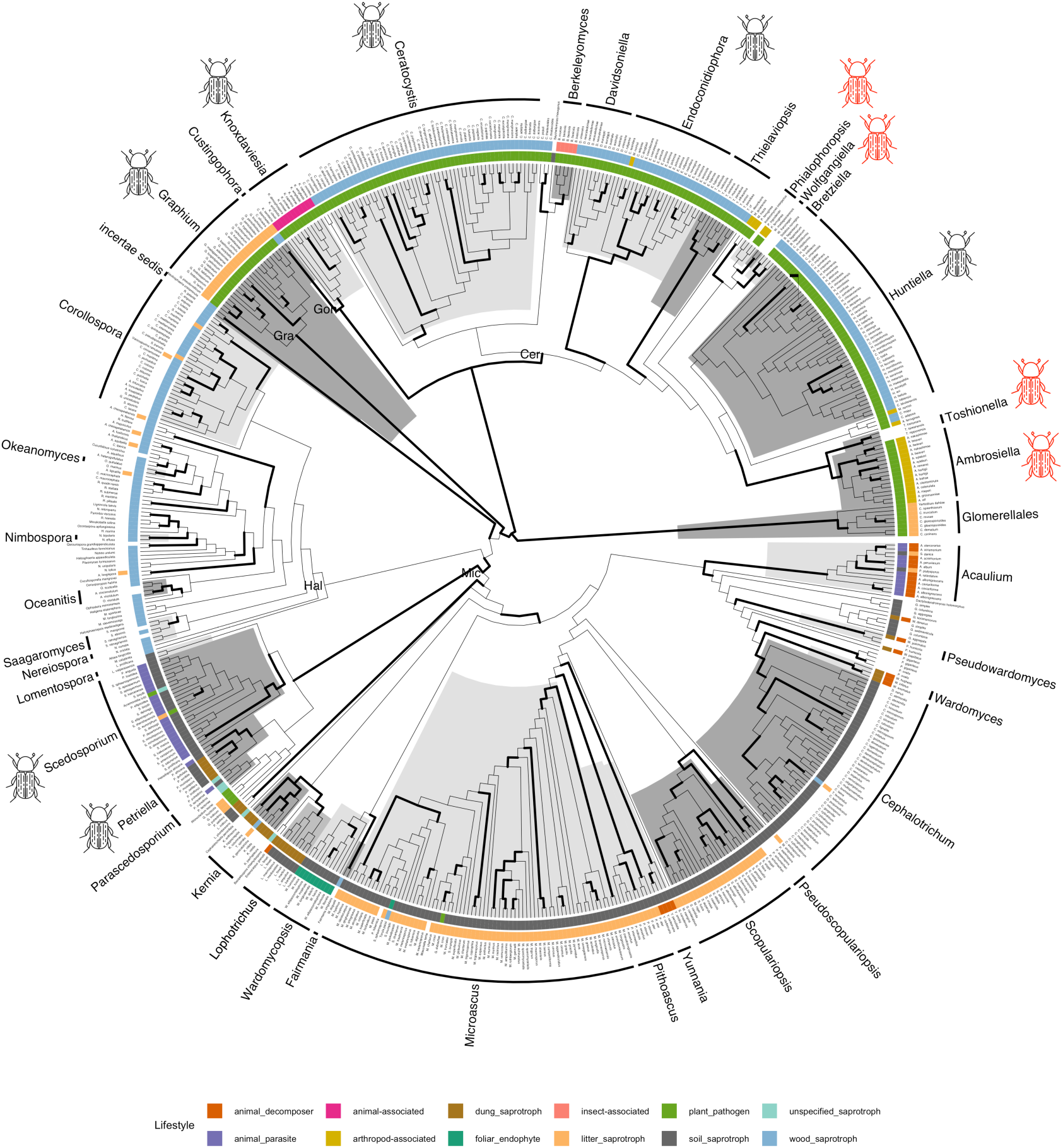
Microascales maximum likelihood species tree, reconstructed using concatenated 7-locus dataset. Bold branches indicate UFBootstrap > 0.95. Higher taxa are are labelled and highlighted by alternating light and dark grey blocks. Coloured tracks show genus-level ecological lifestyle from FungalTraits, inner track is primary lifestyle, outer track is secondary lifestyle. Beetle icons indicate fungal taxa that associate with bark and ambrosia beetles. Red beetle icons indicate beetle-mediated fungal farming lineages i.e. ambrosia fungi. Tree is shown as cladogram for display purpose, see figshare Supplementary Data for branch lengths. Families labelled as follows: Cer = Ceratocystidaceae, Gra = Graphiaceae, Hal = Halosphaeriaceae, Mic = Microascaceae.

The Halosphaeriaceae was paraphyletic with *Nautosphaeria* + *Tubakiella galerita* forming a basal clade to the rest of the family and the Microascaceae. Within the rest of the Halosphaeriaceae, there was widespread polyphyly, with many genera being non-monophyletic (e.g., *Ascosacculus*, *Remispora* s.s., *Aniptodera*, *Magnisphaeria*). The family also contained a high number of monospecific genera.

The Microascaceae was strongly supported, and contained many supported genera, including *Scedosporium*, *Kernia*, *Acaulium*, *Cephalotrichum*, *Wardomyces sensu stricto*, *Pseudoscopulariopsis*, *Scopulariopsis*, *Pithoascus*, *Yunnania*, and a handful of monospecific genera. The genus *Pseudallescheria* was split across various clades, suggesting major revision is needed. *Petriella* and *Petriellopsis* were also unresolved. *Wardomyces sensu lato* also requires revision, as our phylogeny supported *Wardomyces* sensu stricto but not *Parawardomyces* or *Pseudowardomyces*, which both showed a single misplaced taxon. The large genus *Microascus* was monophyletic but only supported by SH-aLRT.

Fungal lifestyles were generally conserved within families but showed clear differences between families (Figure 2). As with the Glomerellales outgroup, all Ceratocystidaceae were reported as plant pathogens, except *Seychellomyces* which was a soil saprotroph. Most Ceratocystidaceae genera were also reported with a secondary wood saprotrophic lifestyle. Insect or arthropod-associated genera in the Ceratocystidaceae (*Ambrosiella*, *Phialophoropsis*, *Meredithiella*, and *Berkeleyomyces*) were scattered across the family and did not form a monophyletic group.

The smaller Gondwanamycetaceae and Graphiaceae families were also primarily plant pathogens, but with differing secondary lifestyles, being animal-associated and litter saprotrophs, respectively. The Halosphaeriaceae stood out as being almost exclusively wood saprotrophic. In contrast, the Microascaceae showed a much more diverse range of lifestyles. Microascaceae genera were mostly saprotrophs, with soil being the most common substrate, followed by dung, ‘unspecified’, and wood. *Acaulium* was an outlier, being primarily an animal parasite. Secondary lifestyles were also variable in the Microascaceae, switching frequently between sister clades among animal parasites, animal decomposers, soil saprotrophs, litter saprotrophs and foliar endophytes.

Bark beetle associations were also non-random. Many Ceratocystidaceae genera were reportedly bark beetle-associated (Figure 2), as well as the smaller Graphiaceae and Gondwanamycetaceae. In contrast, no bark beetle associates were reported for the Halosphaeriaceae, and only one small clade containing the *Scedosporium* and *Petriella* was reported in the Microascaceae. Within the Ceratocystidaceae family, bark beetle-associated genera were not monophyletic. The ambrosia lifestyle was reported in two clades within the Ceratocystidaceae, one containing *Ambrosiella* and *Toshionella* and a second with *Phialophoropsis* and *Wolfgangiella* (Figure 2)

### Ophiostomatales

For the Ophiostomatales, we downloaded 122,424 sequences, of which 15,464 were sorted into the seven loci. Initial sequence filtering left 14,219, and selecting the longest sequence for each taxon at each locus left 3238. After removing unidentified sequences and building initial trees to identify further erroneous sequences, 1853 sequences remained. Concatenating these across loci produced a final concatenated alignment of 748 sequences and 10423 sites, of which 3881 were informative. See Supplementary Table 1 for NCBI accession codes for each locus.

Alignment statistics varied considerably across loci (Table 2). ITS and LSU had the highest number of sequences (495 and 493, respectively), but relatively few informative characters (737 and 332). In contrast, protein-coding genes β-tubulin and EF1a were more balanced, with both high sequence coverage (439 and 360 sequences) and a high number of informative sites (978 and 1209). RPB1 and RPB2 contained fewer sequences (11 and 28, respectively) and showed a high proportion of invariant sites (770 and 651). SSU had 24 sequences and only 63 informative sites.

**Table 2:**
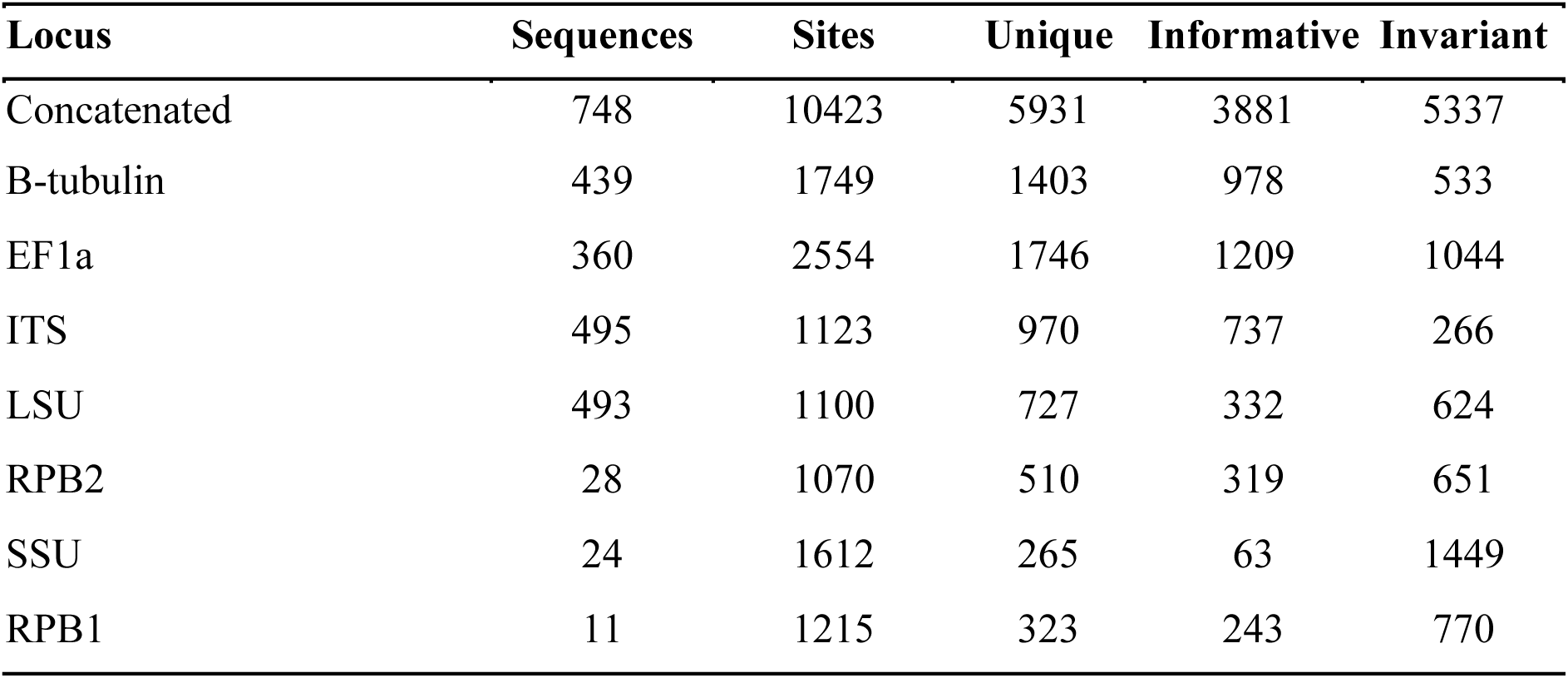
Alignment summary statistics for the Ophiostomatales 7-locus concatenated alignment and individual locus alignments.

| **Locus** | **Sequences** | **Sites** | **Unique** | **Informative** | **Invariant** |
| --- | --- | --- | --- | --- | --- |
| Concatenated | 748 | 10423 | 5931 | 3881 | 5337 |
| B-tubulin | 439 | 1749 | 1403 | 978 | 533 |
| EF1a | 360 | 2554 | 1746 | 1209 | 1044 |
| ITS | 495 | 1123 | 970 | 737 | 266 |
| LSU | 493 | 1100 | 727 | 332 | 624 |
| RPB2 | 28 | 1070 | 510 | 319 | 651 |
| SSU | 24 | 1612 | 265 | 63 | 1449 |
| RPB1 | 11 | 1215 | 323 | 243 | 770 |

Locus-wise assessments of sequence composition and clade support revealed clear differences between loci (Figure 3). As in the Microascales, AT skew varied widely between loci whilst GC% remained relatively stable (Figure 3a and 3b). K2P distance patterns were also very similar to the Microascales apart from RPB2, which showed a noticeably lower K2P in the Ophiostomatales (Figure 3c). A Chi-2 homogeneity of variance test showed nucleotide state frequencies were significantly different between taxa for ITS, β-tubulin, EF1a and RPB2 (*p* < 0.05) (Figure 3f).

**Figure 3:**
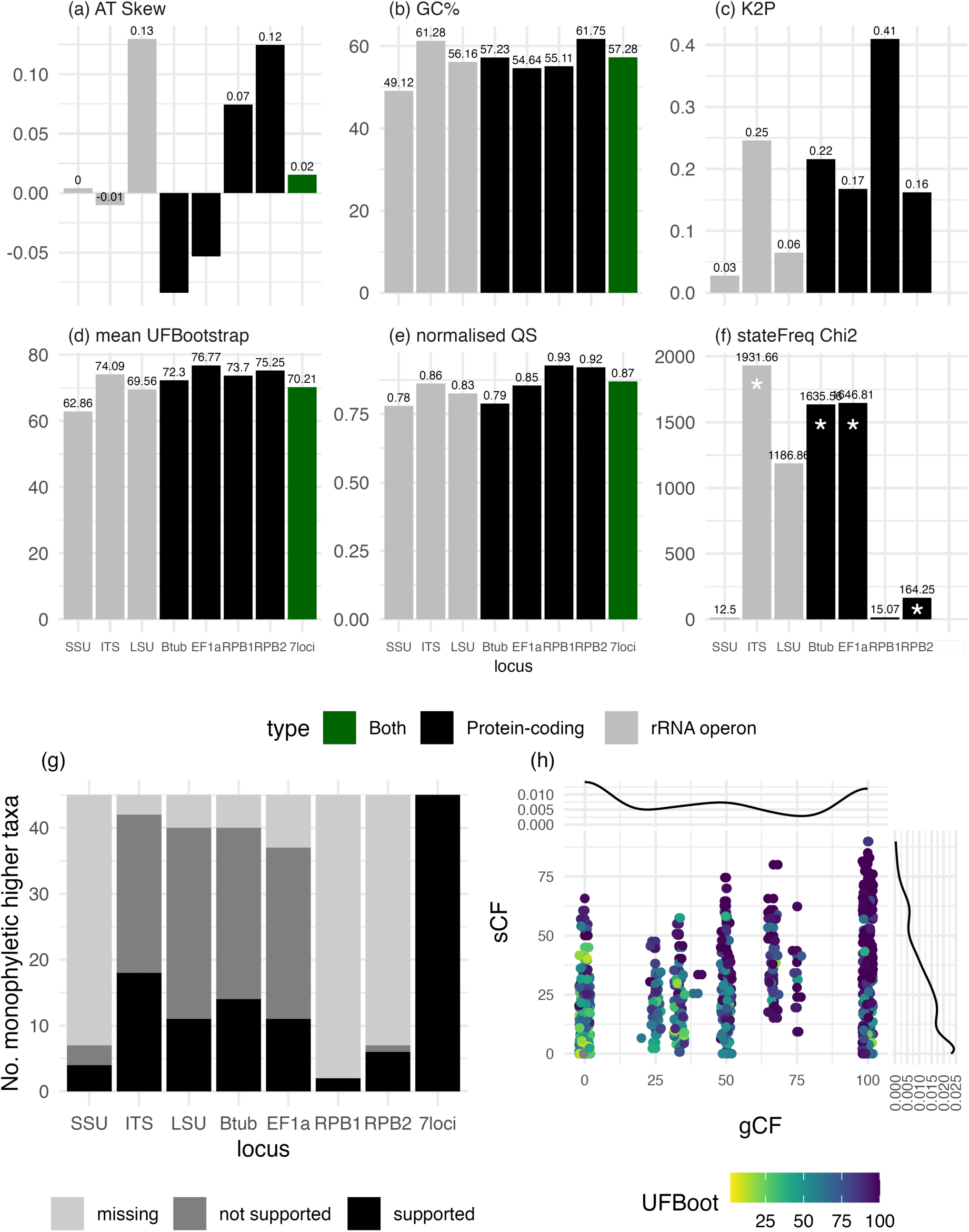
Plots comparing summary statistics of sequence alignments and phylogenetic trees for each locus and the concatenated 7-locus dataset. (a) Alignment AT Skew, (b) alignment GC%, (c) Kimura 2-parameter (K2P) pairwise distance, (d) mean UltraFast Bootstrap support, (e) normalised Quartet Score (QS), (f) nucleotide state frequency Chi-square with asterisk showing significance (*p* < 0.05), (g) number of higher taxa recovered as monophyletic (black), not monophyletic (dark grey), and missing (light grey), (h) scatter plot showing the relationship between IQTree gene Concordance Factor (gCF), site Concordance Factor (sCF), and UFBootstrap. Each point represents one node in the concatenated species tree. Histograms show the distribution of concordance factors.

When comparing locus trees, LSU showed the lowest average bootstrap support at 60.41, whilst RPB1 showed the highest at 91.75 (Figure 3d). Comparing locus trees and the concatenated species tree showed that loci generally agreed with the species tree (QS = 0.87). Individual locus QS did not vary much across loci (Figure 3e). We assessed the number of higher taxa in the species tree that were supported in each locus tree. ITS, LSU, β-tubulin and EF1a all showed similar proportions of supported and unsupported taxa, whilst SSU, RPB1 and RPB2 supported much fewer higher taxa and had a considerably higher number of missing higher taxa (Figure 3g). See Supplementary Table 3 for a detailed list of each higher taxon and its support in each locus tree.

When looking at the concatenated species tree, sCF and gCF showed largely the same pattern as with the Microascales with most nodes receiving low sCF and then a split between very high and very low gCF. In general, nodes with low gCF also tended to have low bootstrap support (Figure 3h)

In the concatenated species tree, the order Ophiostomatales was monophyletic (Figure 4). Within the Ophiostomataceae, *Graphilbum* and *Aureovirgo* formed a basal clade. The basal genera *Heinzbutinia*, *Hawksworthiomyces*, and *Ceratocystiopsis* are well-supported. The rest of the phylogeny was divided into two large clades: one containing the diverse genera *Leptographium* and *Grosmannia*, along with *Raffaelea* sensu stricto, *Dryadomyces*, *Harringtonia*, *Esteya*, and *Fragosphaeria*, and the second containing *Ophiostoma*, *Sporothrix*, and a group of small genera. The *Ophiostoma*, *Spororthix* group was strongly supported.

**Figure 4:**
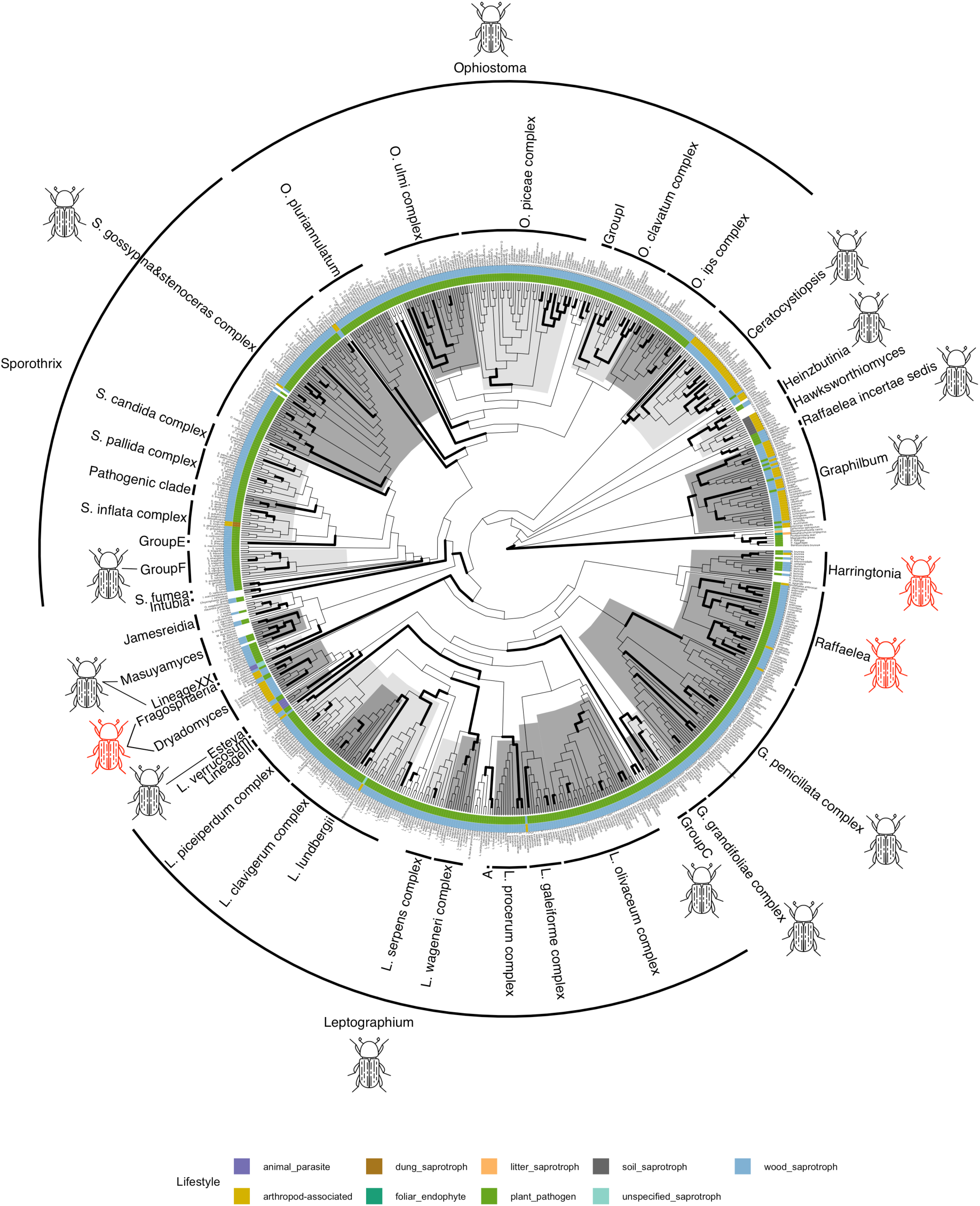
Ophiostomatales Maximum Likelihood species tree, reconstructed using concatenated 7-locus dataset. Bold branches indicate UFBootstrap > 0.95. Higher taxa are are labelled and highlighted by alternating light and dark grey blocks. Coloured tracks show genus-level ecological lifestyle from FungalTraits. Inner track is primary lifestyle, outer track is secondary lifestyle. Beetle icons indicate fungal taxa that associate with bark and ambrosia beetles. Red beetle icons indicate fungal farming lineages i.e. ambrosia fungi. Tree is shown as cladogram for display purpose, see figshare Supplementary Data for tree with branch lengths.

Within the *Leptographium*, *Grosmannia* group, the genera *Fragosphaeria*, *Affroraffaelea*, and *Esteya* formed supported subclades. *Raffaelea* was not monophyletic, with the *R. deltoideospora* group remaining *incertae sedis*. The rest of *Raffaelea sensu stricto* was monophyletic but not supported, although it contained supported subclades. *Grosmannia* was not monophyletic, being split across three species complexes *G. penicillata*, *G. grandifoliae*, and Group C sensu de Beer et al (2022). *Leptographium* was broadly supported, containing various supported species complexes (*L. galeiforme, L. procerum, L. wageneri, and L. clavigerum*). The *L. verrucosum* lineage, however, was placed as sister to *Esteya* and *Dryadomyces*.

Within the *Ophiostoma* and *Sporothrix* clade, *Jamesreidia*, *Chrysosphaeria*, *Intubia* and Lineage XX sensu de Beer et al. (2022) were well supported. *Intubia* and *Chrysosphaeria* formed an unsupported clade subtending the rest of the *Ophiostoma*, *Sporothrix* group. The large genus *Sporothrix* contained many monophyletic species complexes, although most were only supported by SH-aLRT. Within *Ophiostoma*, the pluriannulatum, ulmi, ips, and piceae complexes were recovered, but only the pluriannulatum and piceae complexes received strong support. The average taxonomic retention index (tRI) for the Ophiostomatales species tree was 0.5531.

Fungal lifestyles were highly conserved across the Ophiostomatales, with most genera listed as plant pathogens with a secondary wood saprotrophic lifestyle. The next most common lifestyle was a primary wood saprotroph with a secondary association with arthropods, as seen in *Graphilbum*, *Ceratocystiopsis*, and *Dryadomyces*. Two genera, *Esteya* and *Fragosphaeria,* were reported as animal parasites. The outgroup Magnaporthales showed two additional lifestyles not seen within the Ophiostomatales, foliar endophytes and litter saprotrophs.

Beetle associations were also highly conserved, with almost all genera being reportedly associated with bark and ambrosia beetles. Only the small genera *Hawksworthiomyces*, *Intubia*, *Chrysosphaeria*, *Jamesreidia*, and the *Sporothrix fumea* incertae sedis lineage were not reportedly beetle-associated. Additionally, all species complexes within *Ophiostoma*, *Leptographium*, and *Grosmannia* were beetle-associated. *Sporothrix* was a notable exception, with only the *S. gossypina*/*stenoceras* complex and Group F being beetle-associated. Ambrosia associations were reported in two distantly related clades, one clade with *Harringtonia* and *Raffaelea* and another with *Fragosphaeria* and *Dryadomyces*.

## Discussion

Understanding the evolutionary relationships of ophiostomatoid fungi remains a central challenge, given their often-cryptic lifestyles and the rapid expansion of sequence data. In this study, we present densely sampled phylogenetic trees for the Microascales and Ophiostomatales, integrating all available sequence data for seven loci widely used in fungal molecular taxonomy. This approach allows us to critically evaluate marker choice, sampling depth, and their combined impact on phylogenetic accuracy and taxonomic inference.

### Maximising taxonomic coverage and phylogenetic resolution with locus concatenation and dense, vouchered sampling

When considered individually, marker loci display various weaknesses for phylogenetic analysis, such as varying K2P, and significant differences in state frequencies, and fail to resolve many higher taxa in the Microascales and Ophiostomatales. Loci also differed similarly in both Microascales and Ophiostomatales, suggesting these are true locus differences and not artefacts of the sequences or methods used. However, concatenating loci helps to overcome these issues and significantly improves our ability to robustly resolve phylogenetic relationships in both orders. Furthermore, quartet scores suggest all loci contribute to some extent to the observed species trees, even when taxon sampling was limited.

In the Microascales, ribosomal operon loci ITS and LSU, provided the broadest taxonomic representation but failed to support many genera and species complexes seen in the concatenated species tree (Figure 2). Protein-coding genes (β-tubulin, EF1a, RPB1, RPB2) supported a similar number of genera and higher taxa as ribosomal loci despite sparser taxon sampling, highlighting the need for expanded sampling of these loci. This also reinforces the value of marker concatenation, as it allows us to combine loci with high coverage and those with strong resolution. For the Ophiostomatales, ITS, LSU, β-tubulin, and EF1a showed similar higher taxon support and taxon coverage, suggesting that sampling of protein-coding genes and rRNA loci has been more even than in the Microascales.

Most sequences in this dataset originated from identified, vouchered specimens or culture collections, ensuring traceability and enabling re-examination of morphological and ecological traits. However, compiling and curating sequences into well-sampled, accurate alignments still required considerable manual editing and multiple rounds of tree reconstruction to correct all misidentified, mislabelled or misaligned sequences. Therefore, despite developing a semi-automated pipeline using vouchered material, we still need more efficient approaches for multi-gene phylogenetic studies, especially for diverse fungal lineages such as the Ophiostomatales and Microascales.

### Taxonomic clarifications and novel lineages in the Microascales and Ophiostomatales

Building on this methodological foundation, our phylogenies provide novel insights into the taxonomy of Microascales and Ophiostomatales. In the Microascales, we confirm the subdivision of (Ceratocystidaceae, Gondwanamycetaceae) and (Microascaceae, Graphiaceae, Halosphaeriaceae), suggested by Réblová, Gams and Seifert (2011) and Bao *et al*. (2023).

Most genera within the Ceratocystidaceae and Gondwanamycetaceae are monophyletic, but marine Halosphaeriaceae genera remain problematic. The Microascaceae family also requires revision at the genus level, particularly for *Petriella*, *Pseudallescheria*, and *Wardomyces*.

As in previous studies, *Nautosphaeria* and *Tubakiella* (*Remispora galerita*) form a clade outside current families, supporting recognition of a potential new family within the Microascales (Sakayaroj, Pang and Jones, 2011; Hyde *et al*., 2020; Bao *et al*., 2023). Additionally, we clarify that these genera form a strongly supported clade with Graphiaceae, Microascaceae, and Halosphaeriaceae. However, further work is needed to confirm whether this potential new family is sister to both Halosphaeriaceae and Microascaceae as seen here and in Bao (2023) or whether it’s more closely related to one of two.

In the Ophiostomatales, our results confirm most genera and many species complexes of de Beer *et al*. (2022)’s recent taxonomic revision. Additionally, we recover two large higher clades (*Ophiostoma*, *Sporothrix*, *Intubia*, *Chrysosphaeria, Jamesreidia*, *Masuyamyces*) and (*Dryadomyces, Esteya, Leptographium*, *Grosmannia*, *Raffaelea*, *Harringtonia*) (Figure 4). Subsets of these two clades are consistently recovered in phylogenetic analyses (Nel *et al*., 2021; de Beer *et al*., 2022; Huang *et al*., 2025) and support the idea that the Ophiostomatales may be better split into two smaller, more morphologically distinct families rather than a single large Ophiostomataceae. This was also suggested by de Beer *et al*. (2022), but the authors refrained from doing so due to weak support for deeper nodes on the phylogenetic tree. Here, we clarify for the first time the position of the recently described *Jamesreidia*, *Masuyamyces*, and Lineage XX (sensu de Beer *et al*. (2022)) within one of these two major lineages. However, previous work has also recovered *Fragosphaeria*, *Ceratocystiopsis* and *Hawksworthiomyces* within the *Ophiostoma* clade, which is not seen here.

The placement of *Graphilbum* remains unresolved. Some previous studies resolve it as sister to the *Ophiostoma* + *Sporothrix* clade (Vanderpool, Bracewell and McCutcheon, 2018) and the maximum likelihood tree of Nel *et al*. (2021), whilst others resolve it as sister to the *Leptographium*, *Grosmannia*, *Raffaelea* group (de Beer *et al*., (2022) and the coalescent-based ASTRAL tree of Nel *et al*., (2021)). In our results, it is sister to both groups together with *Aureovirgo* as seen in Huang *et al*. (2025). These drastically different positions suggest that additional data is needed to resolve *Graphilbum*’s position within the family and order.

Another contentious lineage for the Ophiostomatales contains the recently described *Intubia* and *Chrysosphaeria*. Despite lacking strong support, we resolve the two genera as a monophyletic group sister to the *Ophiostoma*, *Sporothrix* clade. This was also seen in Nel *et al*. (2021), who used various tree-building approaches and genome-scale data to clarify their position on the basis that marker genes could not accurately place them. The fact that we were able to recover largely the same Ophiostomatales backbone as genome-based studies (Nel *et al*., 2021; de Beer *et al*., 2022; Huang *et al*., 2025) despite using independent datasets suggests that these marker genes have high phylogenetic information content and that with sufficient taxon sampling, we can still use these marker gene datasets to produce accurate and well-supported phylogenies.

### Fungal lifestyles are conserved in Ophiostomatales but variable in Microascales

The ecological diversity of ophiostomatoid fungi emerges when we annotate the phylogenies with fungal trait data. We observed contrasting patterns between the two orders, with Ophiostomatales generally showing more stable lifestyles than in the Microascales. The same pattern is seen with bark beetle associations, with Ophiostomatales being almost universally beetle-associated, whilst Microascales show much greater variability. Within the Microascales, bark beetle-associated genera are phylogenetically heterogeneous but not randomly drawn from across the whole tree; these genera are concentrated in the diverse Ceratocystidaceae family and smaller Graphiaceae and Gondwanamycetaceae. However, even here they are often sister to non-beetle associates.

These observed differences in fungal lifestyle patterns between orders may partly reflect contrasting approaches to genus delimitation. The Microascales is divided into many small genera, especially within the Halosphaeriaceae, whilst the Ophiostomatales is dominated by a few large, morphologically and ecologically diverse genera. Therefore, genus-level trait data like FungalTraits may look more variable in clades with many small genera like Microascales. Therefore, whilst informative, our broad-scale approach may obscure some important species-level variation.

Displaying beetle-association alongside FungalTrait data also emphasises the flexibility of fungal lifestyles, showing that fungi can have multiple trophic modes whilst facultatively associating with insects. For example, our results reveal a close relationship between beetle association and plant pathogenicity. This reflects a commonly reported mutually beneficial relationship, where beetles and fungi can more effectively overcome host plant defences when colonising plants together (Ploetz *et al*., 2013; Joseph and Keyhani, 2021; Jirošová *et al*., 2022). Retaining their capacity to digest plant compounds for nutrition whilst dispersing efficiently between plants via bark beetles (Davis *et al*., 2019), also explains why these fungi can become such devastating forest pathogens.

It is difficult to infer clear directionality or reconstruct the evolution of specific traits from our trees, as some basal nodes lack strong statistical support. For this reason, we avoided formal trait evolution analyses and instead focused on qualitative interpretations of observed patterns. Still, this dataset presents interesting evolutionary hypotheses for future testing. For example, whilst Ophiostomatales and Microascales form functionally similar beetle symbioses, Microascales associations appear much more plastic, with multiple gains and losses (Mayers *et al*., 2015; Mayers, Harrington, Masuya, *et al*., 2020; Mayers, Harrington, Mcnew, *et al*., 2020). In contrast, the pattern in Ophiostomatales may reflect a more ancient beetle association gained in the common ancestor and then largely maintained with only a few secondary losses in groups such as *Sporothrix*. This stable association in the Ophiostomatales may also explain why specialised fungal domestication and farming evolved so many times independently in groups like *Raffaelea* and *Harringtonia* (Ophiostomatales) (Vanderpool, Bracewell and McCutcheon, 2018).

Our data may also shed some light on the evolution of ambrosia fungi more broadly. Various studies have suggested that Ceratocystidaceae ambrosia lineages are more beetle host specific compared to Ophiostomataceae ambrosia lineages (Skelton *et al*., 2019; Mayers, Harrington, Masuya, *et al*., 2020; Mayers, Harrington and Biedermann, 2022; Osborn *et al*., 2023; Huang *et al*., 2025). If Ophiostomatales ambrosia fungi evolved within a context of pre-existing, widespread beetle association, as suggested here, they may have retained generalist traits resulting in less host specificity. This difference has also been observed at the genomic level, with Ophiostomatales and Microascales showing different genomic adaptations to the ambrosial lifestyle (Huang *et al*., 2025).

## Conclusions

Ultimately, our study demonstrates the value of broad and balanced taxon sampling founded on curated, voucher-derived sequences. We used this data to build a robust phylogenetic framework that we hope can support future research on ophiostomatoid fungi. Accurate, well-sampled phylogenies are particularly useful for interpreting metabarcoding datasets, which are becoming increasingly common for ophiostomatoids. By placing metabarcodes on reference phylogenies rather than reconstructing trees directly from short metabarcode regions, we can measure phylogenetic diversity more reliably and move beyond species or OTU counts to identify field sites with unique or novel lineages. These trees can also accelerate biodiversity exploration, especially in megadiverse or understudied regions, where sequence similarity-based approaches may struggle to confidently assign ophiostomatoid sequences to known taxa. Moreover, by integrating different genetic markers into a unified phylogenetic framework, the same tree can be used to place sequences from different loci—an especially important feature for the Ophiostomatales, which are often difficult to amplify using primers for the universal fungal barcode ITS (Skelton *et al*., 2018). Finally, coupling phylogenetic trees and ecological lifestyle data will allow us to harness the predictive power of phylogenetics and predict traits of unknown samples. Predicting plant pathogenicity of ophiostomatoids, for example, could prove particularly useful for forestry, as it could provide rapid initial screening of the pathogenic potential of fungal isolates, facilitating faster interventions. As sequencing efforts expand and environmental surveys uncover greater species diversity, robust phylogenetic datasets will play an increasingly central role in advancing studies on ophiostomatoid biodiversity, ecology and systematics.

## Acknowledgements

We thank Dr Torda Varga, Emily Hodgson, and Brigid Wong for providing phylogenie scripts and advice on sequence curation. We also thank all the members of the Vogler Lab for their insightful feedback on the datasets presented here. This work was supported by the Leverhulme Trust and carried out for the Leverhulme Centre for the Holobiont (Imperial College London). The funding body had no role in the design of the study, data collection, analysis, interpretation, or writing of the manuscript.

## Data Availability

The phylogenie scripts and multiple sequence alignments and phylogenetic trees generated and analysed during the current study are deposited in the figshare repository https://doi.org/10.6084/m9.figshare.30580976.v1

## Supplementary Materials

**Supplementary Table 1:** Ophiostomatoid sequences used in phylogenetic analyses and associated GenBank accession numbers.

**Supplementary Table 2:**
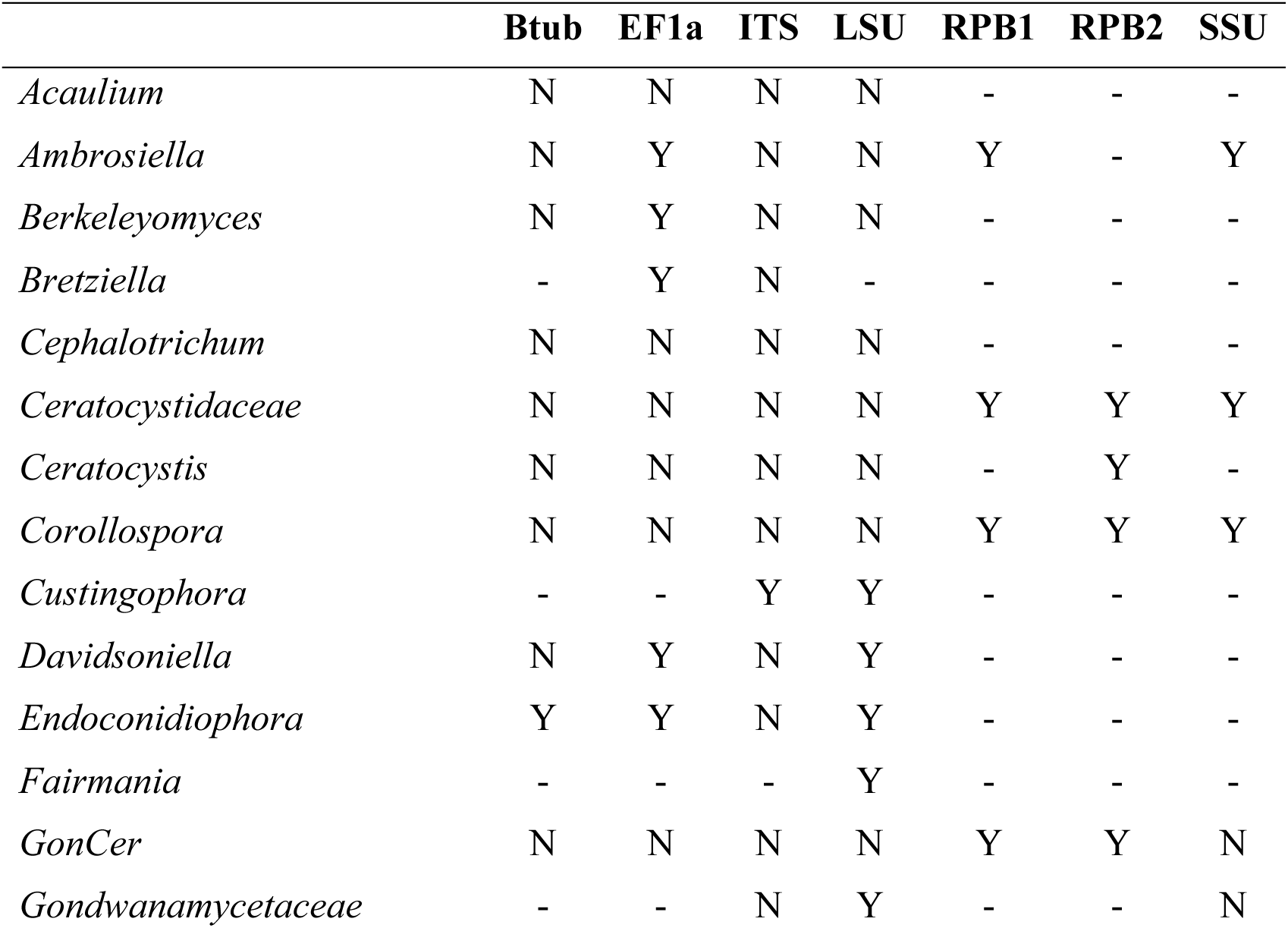
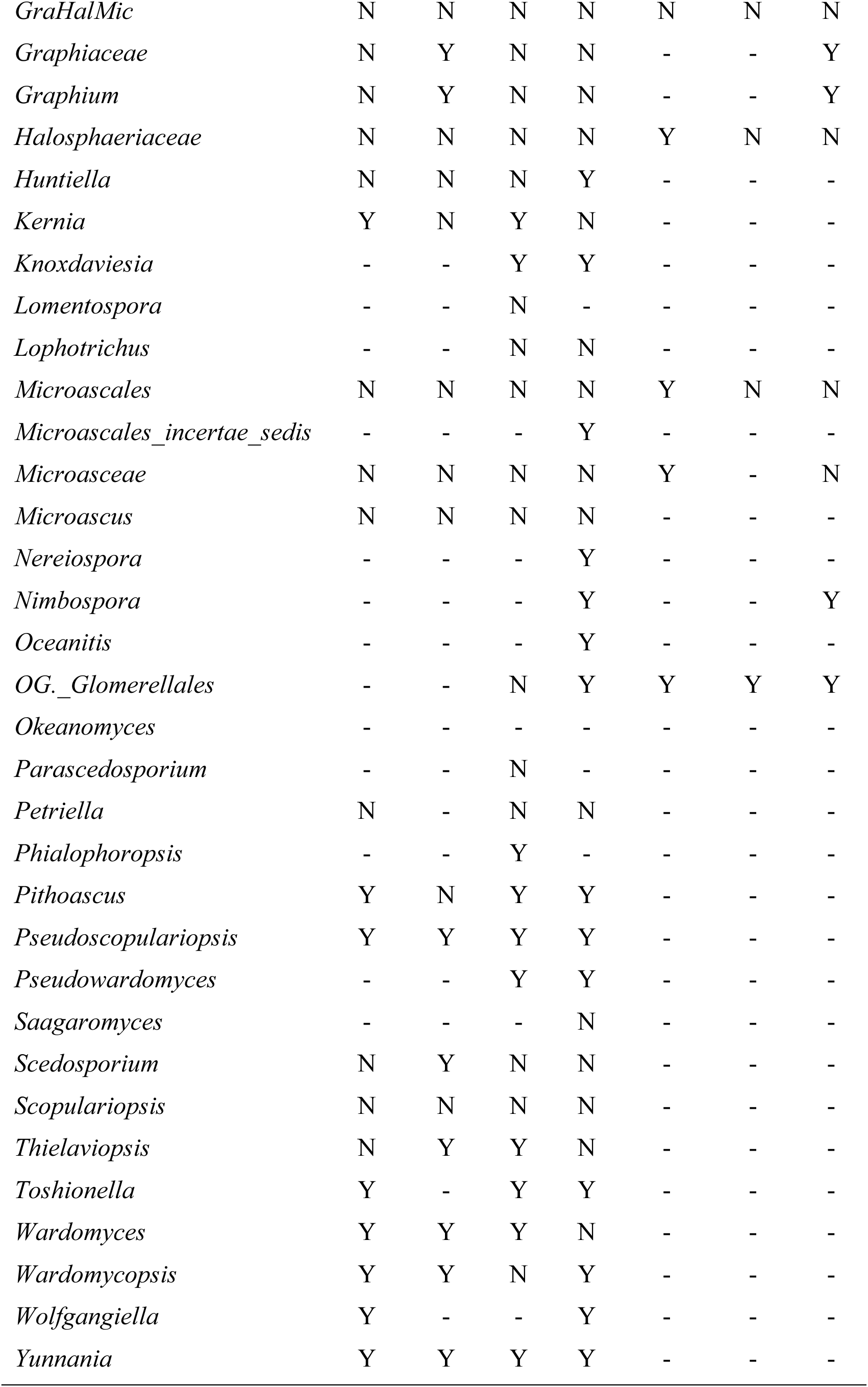
Microascales higher taxa supported by each locus in the 7-locus phylogenetic dataset. Y = supported, N = not supported, ‘-‘ = sequences for that taxon were not present for that locus.

**Supplementary Table 3:**
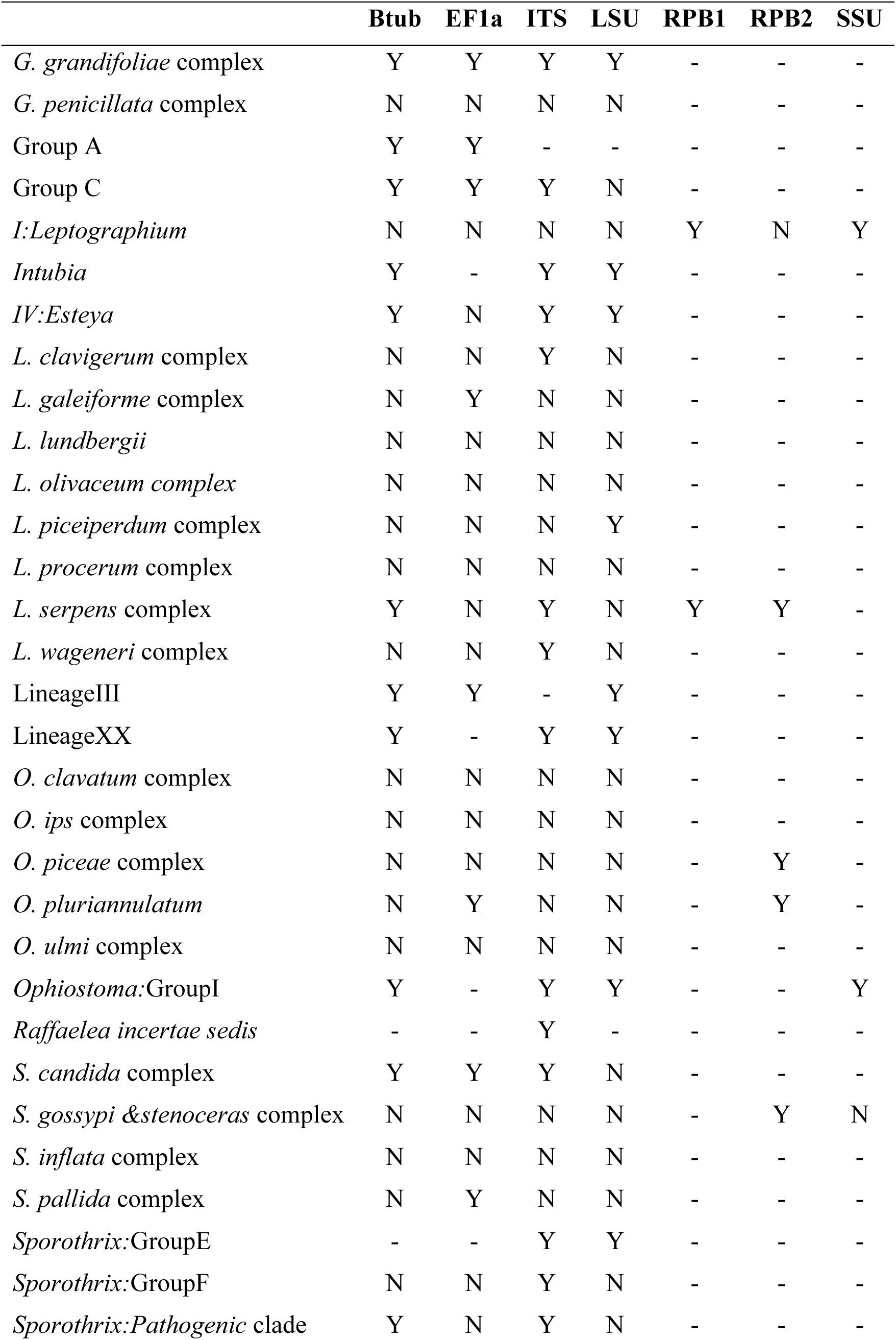
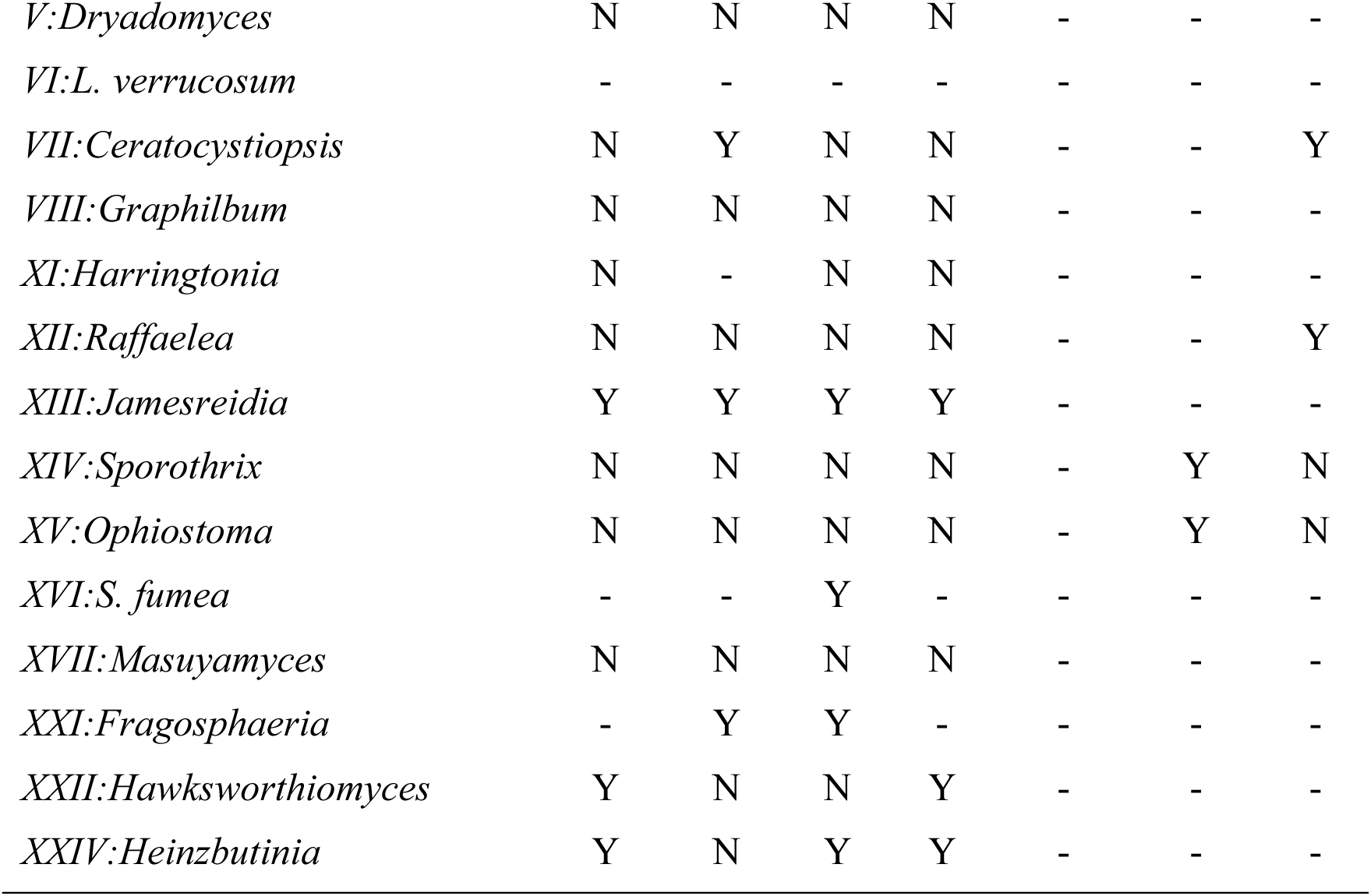
Ophiostomatales higher taxa supported by each locus in the 7-locus phylogenetic dataset. Y = supported, N = not supported, ‘-‘ = sequences for that taxon were not present for that locus. Higher taxon names are formatted to follow the naming of de Beer *et al*. (2022).

**Supplementary Figure 1:**
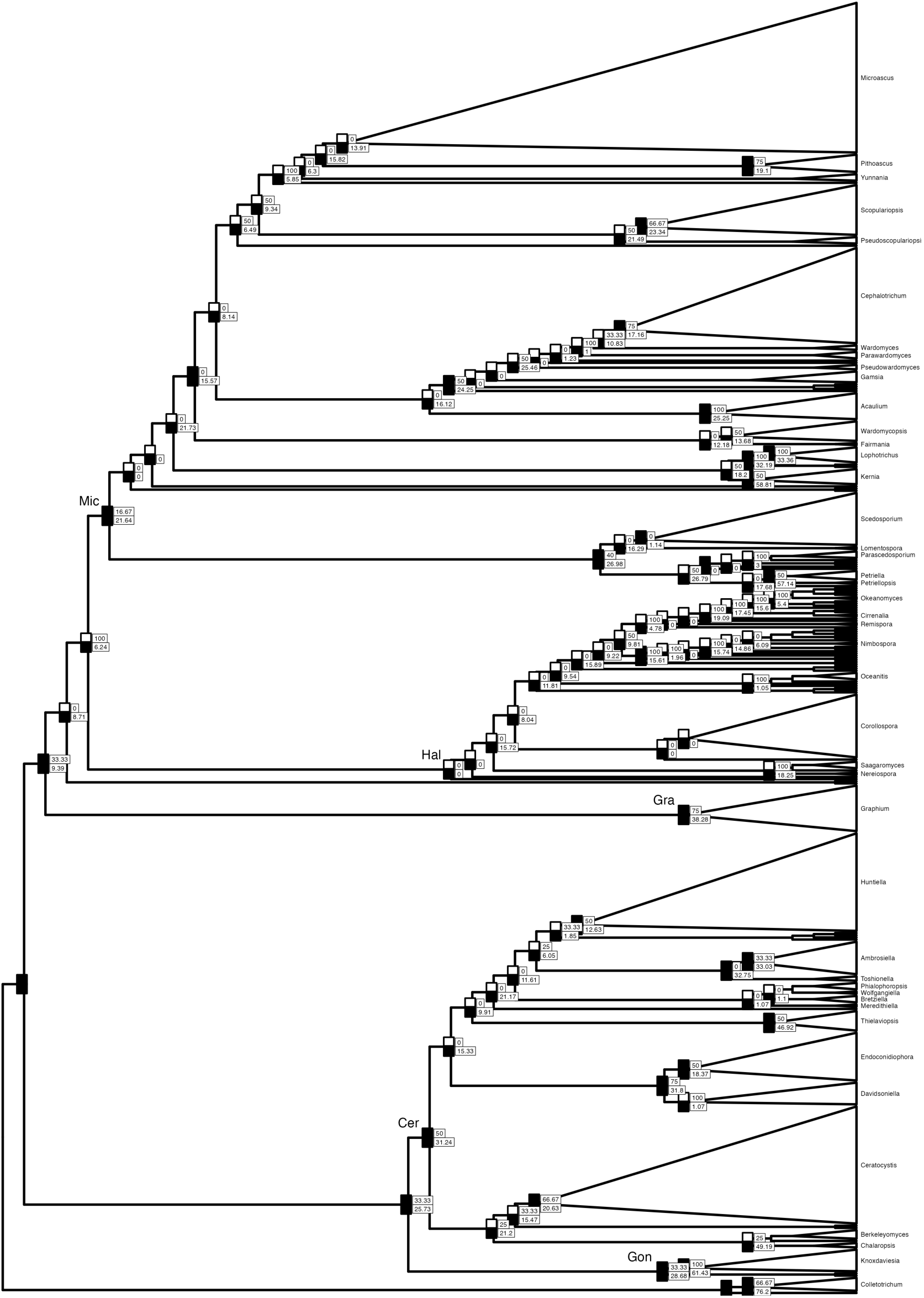
Phylogenetic tree of Microascales shown in Figure Microascales Tree, collapsed to genus level, with node annotations showing UFBootstrap > 0.95 (top left), SH-aLRT > 80 (bottom left), gene concordance factor (top right), site concordance factor (bottom right). Tree is shown as cladogram and was built with IQTree using 7-locus concatenated alignment. Families labelled as follows: Cer = Ceratocystidaceae, Gra = Graphiaceae, Hal = Halosphaeriaceae, Mic = Microascaceae.

**Supplementary Figure 2:**
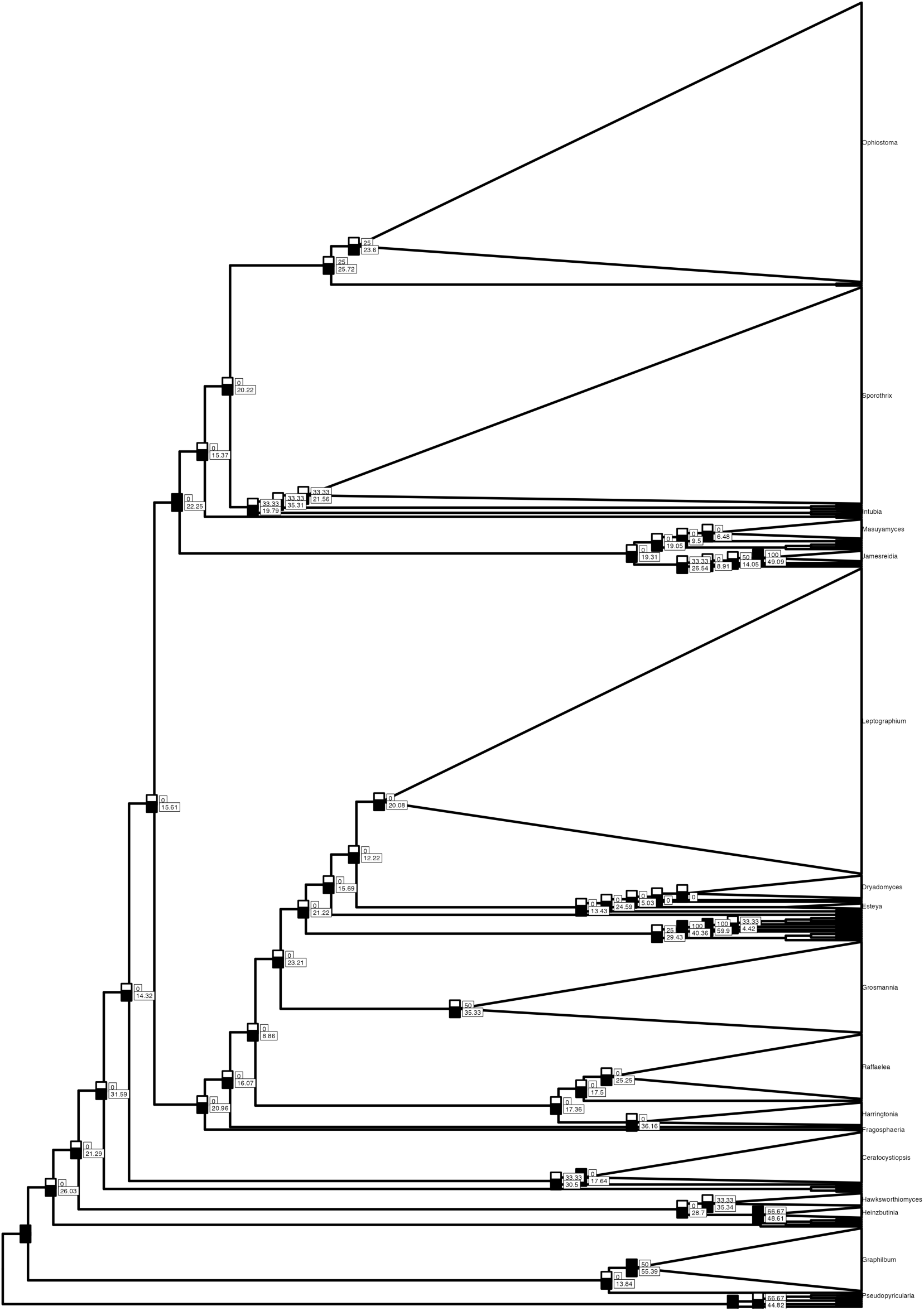
Phylogenetic tree of Ophiostomatales shown in Figure Ophiostomatales Tree, collapsed to genus level, with node annotations showing UFBootstrap > 0.95 (top left), SH-aLRT > 80 (bottom left), gene concordance factor (top right), site concordance factor (bottom right). Tree is shown as cladogram and was built with IQTree using 7-locus concatenated alignment.

